# Replication Study: RAF inhibitors prime wild-type RAF to activate the MAPK pathway and enhance growth

**DOI:** 10.1101/2021.11.30.470372

**Authors:** Steven Pelech, Curtis Gallagher, Catherine Sutter, Lambert Yue, John Kerwin, Ajay Bhargava, Elizabeth Iorns, Rachel Tsui, Alexandria Denis, Nicole Perfito, Timothy M. Errington

**Affiliations:** Kinexus Bioinformatics Corporation, Vancouver, British Columbia, Canada; University of British Columbia, Vancouver, British Columbia, Canada; University of Maryland, College Park, Maryland, United States; NextCure, Beltsville, Maryland, United States; Resilience, Boston, Massachusetts, United States; Shakti BioResearch LLC, Woodbridge, Connecticut, United States; Science Exchange, Palo Alto, California, United States; Komodo Health, San Francisco, California, United States; Center for Open Science, Charlottesville, Virginia, United States; Fordham University School of Law, New York, New York, United States; Rarebase, Palo Alto, California, United States

## Abstract

As part of the Reproducibility Project: Cancer Biology, we published a Registered Report (Bhargava et al., 2016) that described how we intended to replicate selected experiments from the paper “RAF inhibitors prime wild-type RAF to activate the MAPK pathway and enhance growth” (Hatzivassiliou et al., 2010). Here we report the results. We found two unrelated RAF inhibitors, PLX4720 or GDC-0879, selectively inhibited BRAF(V600E) cell proliferation, while the MEK inhibitor, PD0325901, inhibited BRAF(V600E), wild-type RAF/RAS, and mutant RAS cancer cell proliferation, similar to the original study (Figure 1A; Hatzivassiliou et al., 2010). We found knockdown of *CRAF*, but not *BRAF*, in mutant RAS cells attenuated the phospho-MEK induction observed after PLX4720 treatment, similar to the original study (Figure 2B; Hatzivassiliou et al., 2010). The original study reported analogous results with GDC-0879, which was not observed in this replication, although unexpected control results confound the interpretation. We also attempted a replication of an assay with recombinant proteins to test the differential effect of RAF inhibitors on BRAF-CRAF heterodimerization (Figure 4A; Hatzivassiliou et al., 2010). Although we were unable to conduct the experiment as planned, we observed differential binding of BRAF by RAF inhibitors; however, it was between BRAF and beads, independent of CRAF. While these data were unable to address whether, under the conditions of the original study, the same observations could be observed, we discuss key differences between the original study and this replication that are important to consider for further experiments. Finally, where possible, we report meta-analyses for each result.

## Introduction

The <u>Reproducibility Project: Cancer Biology</u> (RP:CB) is a collaboration between the <u>Center for Open Science</u> and <u>Science Exchange</u> that seeks to address concerns about reproducibility in scientific research by conducting replications of selected experiments from a number of high-profile papers in the field of cancer biology (Errington et al., 2014). For each of these papers a Registered Report detailing the proposed experimental designs and protocols for the replications was peer reviewed and published prior to data collection. The present paper is a Replication Study that reports the results of the replication experiments detailed in the Registered Report (Bhargava et al., 2016) for a paper by Hatzivassiliou et al. (2010) and uses a number of approaches to compare the outcomes of the original experiments and the replications.

Conversion of valine-600 to a glutamic amino acid residue (V600E) in the protein kinase encoded by the human BRAF gene is one of the most common mutations in melanomas and many other cancers (Davies et al., 2002), and it produces constitutive activation of its phosphotransferase activity (Kohler and Brummer, 2016). Hatzivassiliou et al. (2010) reported that RAF inhibitors, specifically PLX4720 and GDC-0879, were effective in inhibiting proliferation in BRAF(V600E) cancer cell lines, but not cell lines wild-type for RAF, unlike the MEK inhibitor PD0325901, which inhibited proliferation in both wild-type and BRAF(V600E) cell lines. RAF inhibitor treatment of wild-type RAF cell lines resulted in induction of the MAPK pathway compared to sustained inhibition in BRAF(V600E) cells. Activation of the MAPK pathway by RAF inhibitors in wild-type RAF cells was found to occur in a Raf1 (CRAF) dependent manner (Hatzivassiliou et al., 2010). Priming of CRAF activity was proposed to be mediated by RAF inhibitors’ conformational effects on kinase domain. The CRAF kinase domain and BRAF kinase domain were reported to form a stable complex in the absence of any inhibitor using purified recombinant proteins. Dimerization was destabilized by PLX4720 and stabilized by GDC-0879 with wild-type BRAF. However, the inhibitors had no effect on the interaction between CRAF and BRAF(V600E) kinase domains (Hatzivassiliou et al., 2010). Collectively, these results demonstrated ATP-competitive RAF kinase inhibitors can have opposing functions of the MAPK pathway, depending on cellular context and genotype (Hatzivassiliou et al., 2010), which along with two other reports published at the same time (Heidorn et al., 2010; Poulikakos et al., 2010) provided insight into the mechanism of paradoxical RAF activation in wild-type BRAF cells.

The Registered Report for the paper by Hatzivassiliou et al. (2010) described the experiments to be replicated (Figures 1A, 2B, and 4A), and summarized the current evidence for these findings (Bhargava et al., 2016). The paper by Hatzivassiliou et al. (2010) is one of the first studies that examined the mechanisms underlying the biochemical properties of RAF inhibitors, specifically that while RAF inhibitors suppress RAF activity and downstream MAPK signaling in cells expressing mutant BRAF, these inhibitors do not inhibit and instead paradoxically activate RAF activity and MAPK signaling in cells expressing wild-type BRAF, which has also been reported in other studies (Halaban et al., 2010; Hatzivassiliou et al., 2010; Heidorn et al., 2010; Holderfield et al., 2013; Joseph et al., 2010; Lavoie et al., 2013; Poulikakos et al., 2010). These studies were in agreement that paradoxical activation of MAPK signaling by RAF inhibitors in wild-type BRAF expressing cells required active RAS, but the proposed models differed (Karoulia et al., 2017). Recent studies have continued to elucidate the mechanisms of action of RAF inhibitors. It has been reported that RAF inhibitor binding destabilizes the autoinhibitory interactions that occur between the RAF regulatory and kinase domains, thus promoting the interaction with RAS (Jin et al., 2017; Karoulia et al., 2016). It was also reported that kinase suppressor of Ras 1 (KSR1) is required for the paradoxically activated RAF and MAPK activity induced by RAF inhibitors via drug-induced dimer formation between CRAF and KSR1 (Hu et al., 2011). Additionally, BRAF/CRAF dimer formation in cells treated with RAF inhibitors was demonstrated using a NanoLuc complementation reporter, including differentiating between the reversible property of GDC-0879 and the stable binding of AZ-628 (Dixon et al., 2016). Furthermore, crystal structures of BRAF bound to RAF inhibitors have revealed important insight on the conformation transitions and structural basis that these inhibitors promote dimerization-dependent RAF activation (Karoulia et al., 2016; Thevakumaran et al., 2015). Thus, RAF inhibitors, particularly new inhibitors currently under preclinical and clinical development, have been broadly classified into two main classes, ‘αC-IN’ or ‘αC-OUT’, according to the confirmation in which they stabilize their target kinase (Durrant and Morrison, 2018; Karoulia et al., 2017).

**Figure 1.**
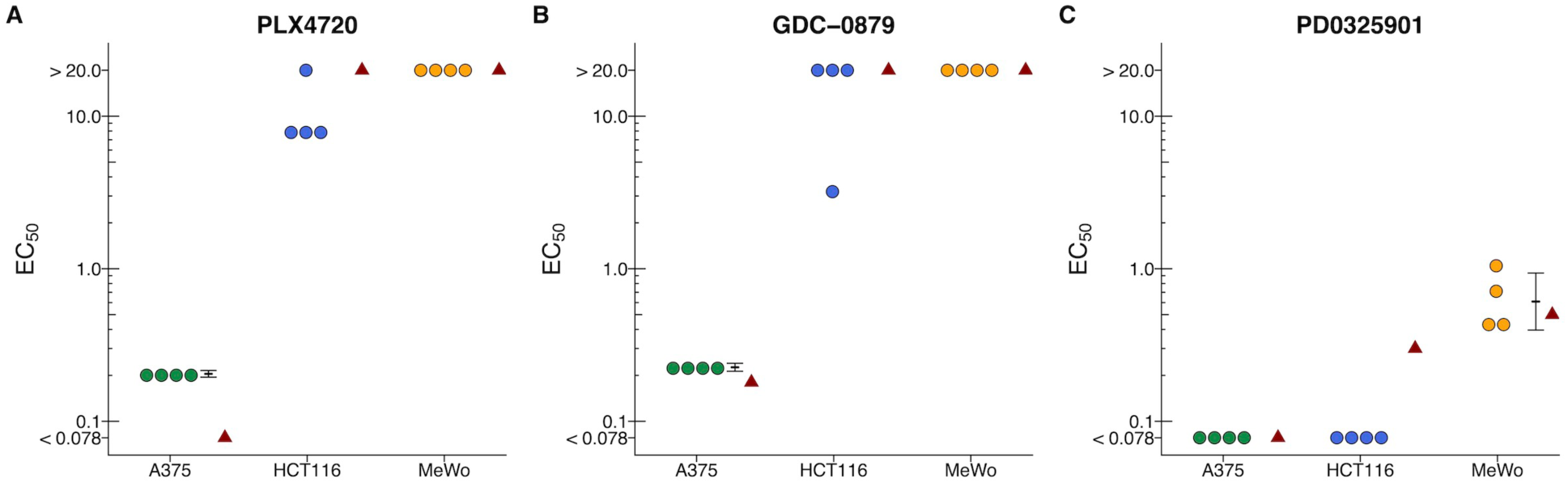
Cellular dose response curves for RAF and MEK inhibitors in BRAF(V600E), mutant RAS, and wild-type RAF/RAS cell lines. Cell viability assays were performed for RAF inhibitors (PLX4720 and GDC-0879) and MEK inhibitor (PD0325901) against A375, (BRAF(V600E)), HCT116 (mutant RAS), and MeWo (wild-type RAF/RAS) cells. Absolute half-maximum effective concentration (EC_50_) values (µM) were determined for each biological repeat. EC_50_ values unable to be accurately estimated are reported as either >20 µM or <0.078 µM, which are the largest and smallest doses tested, respectively. Absolute EC_50_ values for each biological repeat for A375, HCT116, and MeWo cells treated with (**A**) PLX4720, (**B**) GDC-0879, or (**C**) PD0325901 for this replication attempt are plotted with the EC_50_ value reported in Hatzivassiliou et al. (2010) displayed as a single point (red triangle) for comparison. Where possible the mean and 95% confidence interval of the replication data are shown. Additional details for this experiment can be found at https://osf.io/52zp9/.

**Figure 2.**
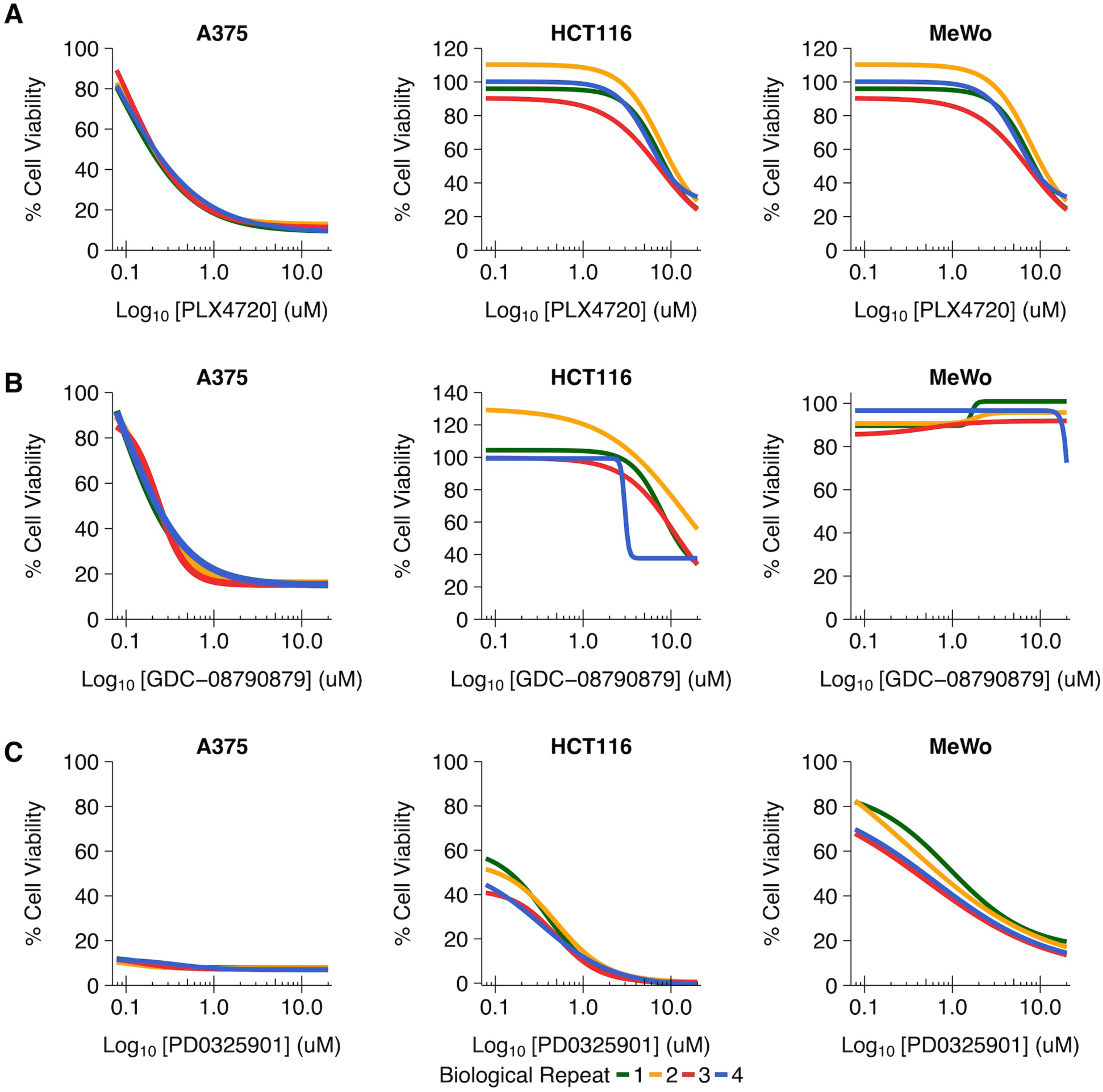
Cellular dose response curves for each biological repeat. This is the same experiment as in Figure 1. The dose response curve of each biological repeat [n=4] for A375, HCT116, and MeWo cells treated with (**A**) PLX4720, (**B**) GDC-0879, or (**C**) PD0325901 for this replication attempt are plotted. Additional details for this experiment can be found at https://osf.io/52zp9/.

The outcome measures reported in this Replication Study will be aggregated with those from the other Replication Studies to create a dataset that will be examined to provide evidence about reproducibility of cancer biology research, and to identify factors that influence reproducibility more generally.

## Results and Discussion

### Assessing cell viability of a panel of cancer cell lines treated with RAF and MEK inhibitors

Using the same ATP-competitive RAF inhibitors as Hatzivassiliou et al. (2010), we aimed to independently replicate an experiment testing the selective nature of RAF inhibitors in inhibiting proliferation of BRAF(V600E) cells. Similar to the original study, cancer cells were treated with various doses of the RAF inhibitors, PLX4720 or GDC-0879, or the MEK inhibitor PD0325901, and cell viability was determined four days later. This experiment was comparable to what was reported in Figure 1A of Hatzivassiliou et al. (2010) and described in Protocol 1 in the Registered Report (Bhargava et al., 2016). While the original study included a panel of BRAF(V600E), wild-type RAF/RAS, and mutant RAS cell lines, this replication attempt was restricted to one cell line from each model. For each cell line, half-maximum effective concentration (EC_50_) values were determined for each inhibitor (Figure 1, Figure 2). The RAF inhibitors, PLX4720 and GDC-0879, inhibited proliferation in A375 cells, which feature BRAF(V600E), with a mean EC_50_ of 0.20 µM, 95% CI [0.19, 0.22] and 0.23 µM, 95% CI [0.21, 0.24], respectively. In contrast, HCT116 cells, which feature a RAS mutant, and MeWo cells, which are wild-type for RAF and RAS, the EC_50_ values for both RAF inhibitors were either more than a log_10_ increase over the EC_50_ values from A375 cells or unable to be accurately determined following published guidelines (Sebaugh, 2010) and thus depicted as > 20 µM, the highest dose tested (Figure 1A-B). However, the MEK inhibitor PD0325901 inhibited proliferation of all cell lines with an EC_50_ value for A375 and HCT116 cells that was unable to be accurately determined and instead depicted as < 0.078 µM, the lowest dose tested (Figure 1C), and MeWo cells resulting in a mean EC_50_ of 0.61 µM, 95% CI [0.40, 0.94]. The original study reported inhibition of cellular proliferation for A375 cells with an estimated EC_50_ of 0.5 µM for PLX4720 and 0.3 µM for GDC-0879, while EC_50_ values for HCT116 and MeWo cells were unable to be determined (> 20 µM) for both RAF inhibitors (Hatzivassiliou et al., 2010). In the original study, the MEK inhibitor inhibited cellular proliferation of all cell lines with an estimated EC_50_ value of 0.18 µM for HCT116 cells and undetermined values for A375 and MeWo cells [< 0.078 µM]. To summarize, for this experiment we found results that were in the same direction as the original study.

### Assessing CRAF and BRAF roles in drug-dependent activation of MEK

To test the role that CRAF and BRAF have in signalling to MEK in non-BRAF(V600E) cells after RAF inhibitor treatment HCT116 cells engineered to stably express dox inducible shRNA (Hoeflich et al., 2006) were utilized. This experiment is similar to what was reported in Figure 2B of Hatzivassiliou et al. (2010) and described in Protocol 2 in the Registered Report (Bhargava et al., 2016). Cells were treated for one hour with varying concentrations of PLX4720 or GDC-0879 and then analyzed for phospho-MEK levels. For cells treated with PLX4720, the induced knockdown efficiency of BRAF or CRAF was on average 80% and 95%, respectively (Figure 3C,D). When CRAF was depleted, an attenuated induction of phospho-MEK was observed, but not when BRAF was depleted (Figure 3A,B). For comparison, the original study reported knockdown of CRAF, but not BRAF, reversed the phospho-MEK induction after PLX4720 treatment. Interestingly, the pattern of induction in the original study, with a dose-dependent increase in phospho-MEK, was not observed in this replication attempt, which instead consistently observed a peak induction with the lowest dose tested (Figure 3A,B).

**Figure 3.**
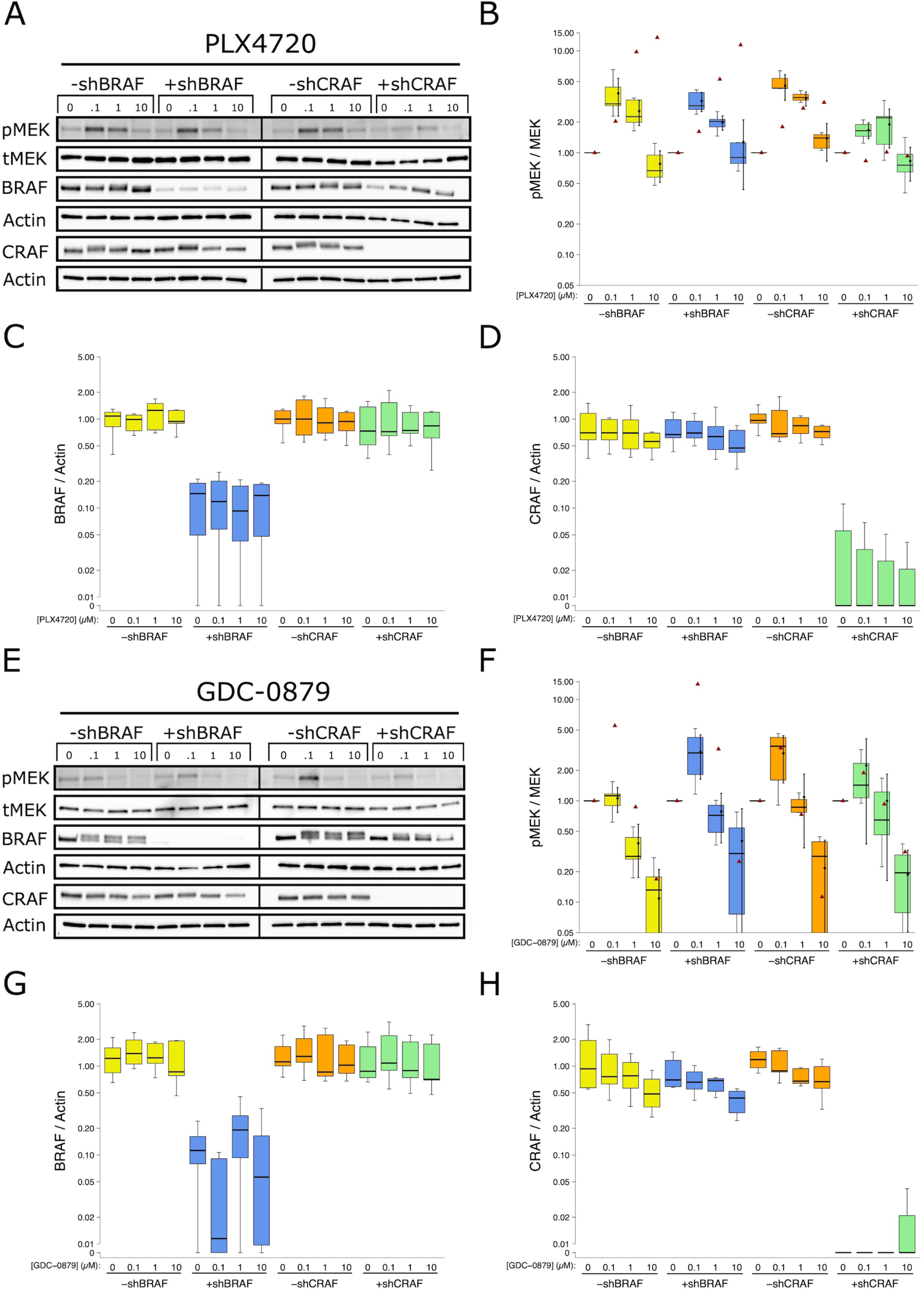
Role of BRAF and CRAF in non-BRAF(V600E) RAF inhibitor induced MEK phosphorylation. HCT116 (mutant RAS) cells expressing doxycycline-inducible shRNA directed against *BRAF* or *CRAF* were treated with RAF inhibitors (PLX4720 or GDC-0879). (**A**) Representative Western blots (experiment performed independently 7 times) of lysates from cells treated with DMSO or the indicated doses of PLX4720 for 1 hr in the absence (-shRNA) or presence (+shRNA) of doxycycline. Membranes were probed with phospho-MEK (pMEK), total MEK (tMEK), BRAF (Actin served as loading control), and CRAF (Actin served as loading control) specific antibodies. (**B**) Western blot bands were quantified for cells treated with PLX4720 and for each biological repeat [n=7] pMEK levels were normalized to tMEK and then to DMSO for each condition. Box and whisker plot with median represented as the line through the box and whiskers representing values within 1.5 IQR of the first and third quartile. Means as block dot and bold error bars represent 95% CI. Data estimated from the representative experiment reported in Figure 2B of Hatzivassiliou et al., 2010 is displayed as a single point (red triangle) for comparison. Statistical analysis was performed on natural log transformed data generated during this replication attempt. Two-way ANOVA interaction between shRNA (*BRAF* and *CRAF*) and induction (no dox or dox); *F*(1,80) = 6.95, *p* = 0.010. Pairwise contrast between -shBRAF and +shBRAF; *t*(80) = 0.071, uncorrected *p* = 0.94, corrected *p* > 0.99, Cohen’s *d* = -0.02, 95% CI [-0.63, 0.58]. Pairwise contrast between -shCRAF and +shCRAF; *t*(80) = 3.66, uncorrected *p* = 0.00045, corrected *p* = 0.00091, Cohen’s *d* = 1.13, 95% CI [0.47, 1.78]. (**C**) Box and whisker plot of quantified Western blot bands for cells treated with PLX4720. BRAF levels from each biological repeat normalized to Actin. (**D**) Box and whisker plot of quantified Western blot bands for cells treated with PLX4720. CRAF levels from each biological repeat normalized to Actin. (**E**) Representative Western blots (experiment performed independently 7 times) of lysates from cells treated with DMSO or the indicated doses of GDC-0879 for 1 hr in the absence (-shRNA) or presence (+shRNA) of doxycycline. Membranes were probed with same antibodies described in **A**. (**F**) Western blot bands were quantified for cells treated with GDC-0879 and normalized as described in **B** for each biological repeat [n=7]. Box and whisker plot shown. Means as block dot and bold error bars represent 95% CI. Data estimated from the representative experiment reported in Figure 2B of Hatzivassiliou et al. (2010) is displayed as a single point (red triangle) for comparison. Statistical analysis was performed on natural log transformed data generated during this replication attempt. Comparison of -shBRAF and +shBRAF; *U* = 131, uncorrected *p* = 0.024, corrected *p* = 0.048, Cliff’s delta = -0.41, 95% CI [-0.65, -0.08]. Comparison of -shCRAF and +shCRAF; *U* = 247, uncorrected *p* = 0.505, corrected *p* > 0.99, Cliff’s delta = 0.12, 95% CI [-0.21, 0.43]. (**G**) Box and whisker plot of quantified Western blot bands for cells treated with GDC-0879. BRAF levels from each biological repeat normalized to Actin. (**H**) Box and whisker plot of quantified Western blot bands for cells treated with GDC-0879. CRAF levels from each biological repeat normalized to Actin. Additional details for this experiment can be found at https://osf.io/dqr5d/.

To determine if the observed results were statistically significant, the analysis plan specified in the Registered Report (Bhargava et al., 2016) proposed to compare the normalized phospho-MEK levels from the *BRAF* or *CRAF* shRNA cell lines (with or without dox induction) across the varying doses of PLX4720. Importantly, while there are various ways to normalize the phospho-MEK levels, the strategy to normalize phospho-MEK to total MEK was specified prior to data collection and analysis to minimize confirmation bias (Wagenmakers et al., 2012). To test if induction of *CRAF* shRNA, but not *BRAF* shRNA, was effective in reducing phospho-MEK levels, we performed an analysis of variance (ANOVA) having two levels of shRNA (*CRAF* or *BRAF*) and two levels of induction (no dox or dox). The ANOVA result on phospho-MEK levels from PLX4720 treated cells normalized to vehicle control (natural log-transformed) was statistically significant for the interaction effect, *F*(1,80) = 6.95, *p* = 0.010. Thus, the null hypothesis that induction of *CRAF* shRNA and induction of *BRAF* shRNA have similar effects on phospho-MEK induction can be rejected. The main effect for dox induction, *F*(1,80) = 6.43, *p* = 0.013 was also statistically significant, while the main effect for shRNA, *F*(1,80) = 0.007, *p* = 0.933 was not. As outlined in the Registered Report (Bhargava et al., 2016), we planned to conduct two comparisons using the Bonferroni correction to adjust for multiple comparisons, making the *a priori* adjusted significance threshold 0.025. The planned comparison of the normalized phospho-MEK levels from *BRAF* shRNA expressing cells without dox induction compared to cells with dox induction was not statistically significant (*t*(80) = 0.071, uncorrected *p* = 0.94, corrected *p* > 0.99, Cohen’s *d* = -0.02, 95% CI [-0.63, 0.58]), while the planned comparison from *CRAF* shRNA expressing cells without dox induction compared to cells with dox induction was statistically significant (*t*(80) = 3.66, uncorrected *p* = 0.00045, corrected *p* = 0.00091, Cohen’s *d* = 1.13, 95% CI [0.47, 1.78]). Thus, the null hypothesis that expression of *CRAF* shRNA has similar effects on phospho-MEK after PLX4720 treatment as cells that do not express the *CRAF* shRNA can be rejected. To summarize, for this experiment we found results that were in the same direction as the original study and statistically significant where predicted.

The role of CRAF and BRAF was also tested in the same experimental set-up with GDC-0879. The induced knockdown efficiency of BRAF or CRAF was on average 91% or 85%, respectively (Figure 3G,H). Unlike PLX4720, there was little to no change in induction of phospho-MEK observed when BRAF or CRAF was depleted in GDC-0879 treated cells (Figure 3E,F). For comparison, the original study reported depletion of CRAF, but not BRAF, reversed the phospho-MEK induction after GDC-0879 treatment, similar to what was reported for PLX4720. Interestingly, in this replication attempt there was little to no induction of phospho-MEK by GDC-0879 treatment in the cells not induced to express the *BRAF* shRNA (-shBRAF condition in Figure 3A,B), even though the increase in phospho-MEK was observed in the other conditions. We also observed a mobility shift in BRAF with GDC-0879 treatment (Figure 3E). This is similar to the mobility shift observed after treatment with the RAF inhibitors sorafenib and 885-A, or PLX4720 co-treated with the MEK inhibitor PD184352, that was previously suggested to be due to hyperphosphorylation of BRAF by MEK-ERK-dependent and MEK-ERK-independent mechanisms (Heidorn et al., 2010).

Similar to the PLX4720 data, the analysis plan specified in the Registered Report (Bhargava et al., 2016) proposed to compare the normalized phospho-MEK levels from the *BRAF* or *CRAF* shRNA cell lines (with or without dox induction) across the varying doses of GDC-0879 with two planned comparisons. Since the data were not normally distributed, Wilcoxon-Mann-Whitney tests (non-parametric equivalent of a *t*-test) were performed using the Bonferroni correction to adjust for multiple comparisons, making the *a priori* adjusted significance threshold 0.025. The planned comparisons of the normalized phospho-MEK levels from *BRAF* shRNA without dox induction compared to cells with dox induction was statistically significant (*U* = 131, uncorrected *p* = 0.024, corrected *p* = 0.048, Cliff’s delta = -0.41, 95% CI [-0.65, -0.08]) and the comparison of *CRAF* shRNA expression cells was not (*U* = 247, uncorrected *p* = 0.505, corrected *p* > 0.99, Cliff’s delta = 0.12, 95% CI [-0.21, 0.43]). To summarize, for this experiment we found results that were not in the same direction as the original study and not statistically significant where predicted.

Interpretation of these results should take into consideration that GDC-0879 did not appreciably activate MEK in the *BRAF* shRNA cell line (Figure 3E-F; -shBRAF lanes), even though this occurred in the *CRAF* shRNA cell line (Figure 3E-F; -shCRAF lanes). This was observed with all replicates (Figure 3F), indicating that this was systemic to the overall experimental design and execution. Additionally, this was not the case for PLX4720, which displayed similar activation of MEK in the *BRAF* and *CRAF* shRNA cells, albeit with unexpected suppression of pMEK at high concentrations. An important aspect of the experimental design that might explain this is that the *BRAF* and *CRAF* shRNA cells, although derived from the same parental HCT116 cell lines, were generated independently to express the indicated dox-inducible shRNA (Hoeflich et al., 2006). Thus, it is possible that while the shRNA is maintained, as indicated by the consistent knockdown of CRAF or BRAF (Figure 3), the response to GDC-0879 under conditions that do not express the shRNAs differ between the cells as a result of genetic drift during maintenance of the cultures, particularly the *BRAF* shRNA cells. This could also be due to differences in growth conditions between laboratories, as well as the inherent variability of the Western blot technique, especially with the use of chemiluminescence, an enzymatic-based technique affected by a number of factors (Koller and Wätzig, 2005).

### Biochemical heterodimerization assay with recombinant RAF proteins in the presence or absence of RAF inhibitors

Using purified recombinant proteins, we aimed to replicate whether the CRAF kinase domain forms a stable complex with the BRAF kinase domain and if the ATP-competitive RAF inhibitors modulate this dimerization. This experiment tested whether RAF priming is mediated by the inhibitors’ conformational effects on the RAF kinase domain and is similar to what was reported in Figure 4A of Hatzivassiliou et al. (2010) and described in Protocol 3 in the Registered Report (Bhargava et al., 2016). The original study reported that the interaction between the CRAF and BRAF kinase domains was destabilized in the presence of the RAF inhibitor PLX4720 or AMP-PCP (a non-hydrolysable analogue of ATP), whereas in the presence of the RAF inhibitor GDC-0879 the dimerization was stabilized. By contrast, these compounds failed to alter the basal interactions between the CRAF and BRAF(V600E) domains. Although not included in this replication attempt, the original study also included the chemically unrelated ATP-competitive RAF inhibitor AZ-628. The dimerization assay was performed similar to the original study. That is, after incubation of the purified recombinant proteins with or without the RAF inhibitors, an immunoprecipitation (IP) assay was performed using the recombinant GST-tagged CRAF kinase domain as the bait and either the wild-type or BRAF(V600E) kinase domain as the putative prey. After the IP was performed, using anti-GST antibodies bound to Protein-A-agarose beads, the bound proteins were detected via Western blot using BRAF-and CRAF-specific antibodies.

**Figure 4.**
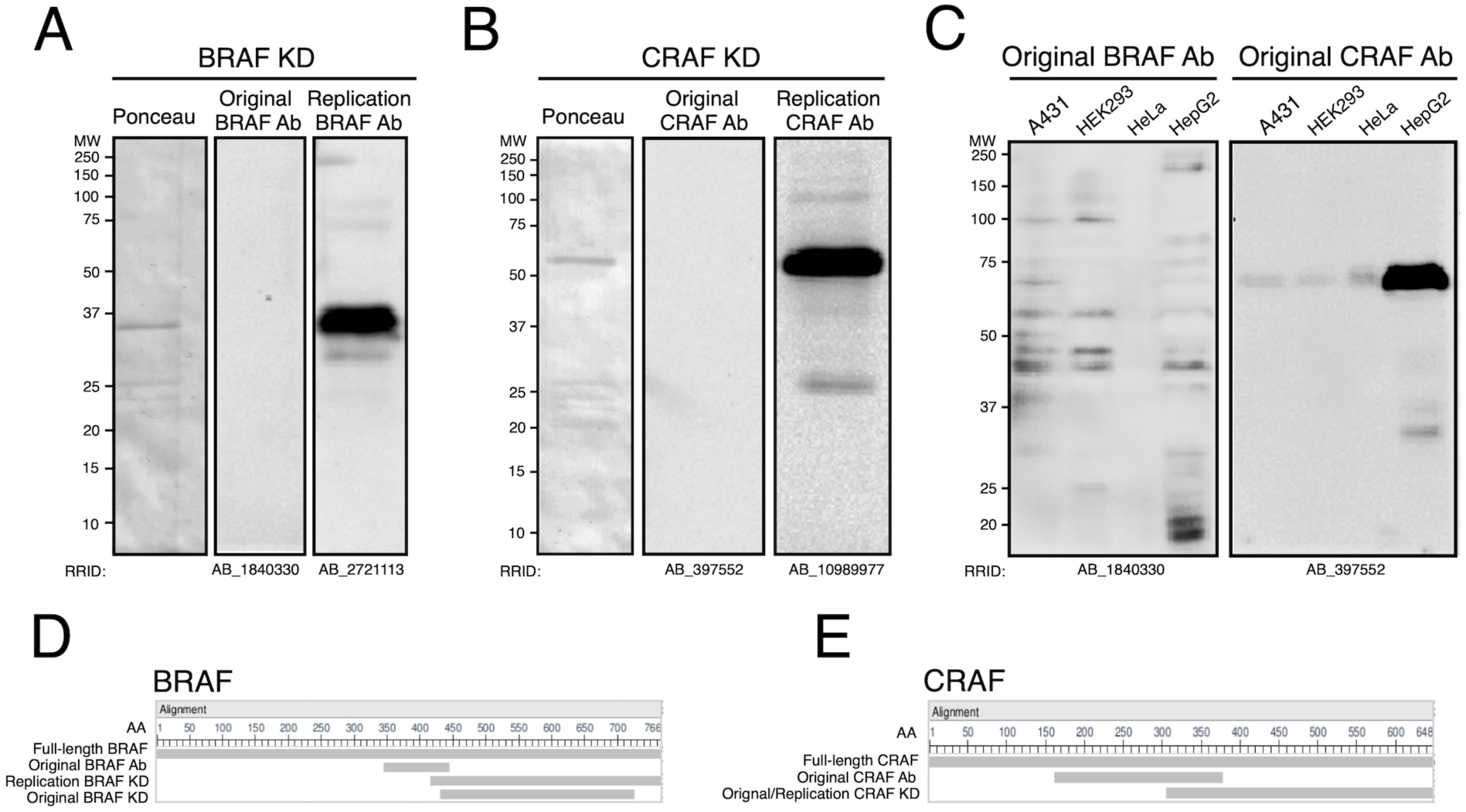
Testing of BRAF and CRAF antibodies used in original study. The BRAF and CRAF antibodies reported in the original study (Hatzivassiliou et al., 2010) were tested for suitability by Western blot. (**A**) Representative Western blots and Ponceau-stained membrane (experiment performed independently 3 times) of 0.5 µg of purified recombinant BRAF kinase domain (KD). Membranes were probed with the BRAF antibody used in the original study (RRID:AB_1840330) or with a different BRAF antibody used in this replication attempt (RRID:AB_2721113). (**B**) Representative Western blots and Ponceau-stained membrane (experiment performed independently 3 times) of 0.5 µg of purified recombinant GST tagged CRAF KD. Membranes were probed with the CRAF antibody used in the original study (RRID:AB_397552) or with a different CRAF antibody used in this replication attempt (RRID:AB_10989977). (**C**) Representative Western blots (experiment performed once) of 16 µg of whole cell lysates from the indicated human cell lines. Membranes were probed with the BRAF (RRID:AB_1840330) or CRAF (RRID:AB_397552) antibodies used in the original study. (**D**) Screenshot from the Constraint-based Multiple Alignment Tool (COBALT) (Papadopoulos and Agarwala, 2007) showing the alignment of full-length BRAF (NCBI Reference Sequence: <u>NP_004324.2</u>), the region of BRAF used as the immunogenic peptide used to generate the BRAF antibody used in the original study (RRID:AB_1840330), the BRAF KD used in this replication attempt, and the BRAF KD used in the original study. (**E**) Screenshot from COBALT showing the alignment of full-length CRAF (NCBI Reference Sequence: <u>NP_002871.1</u>), the region of CRAF used as the immunogenic peptide used to generate the CRAF antibody used in the original study (RRID:AB_397552), and the CRAF KD used in the original study and this replication attempt. Additional details for this experiment can be found at https://osf.io/6f3sk/.

We first performed initial attempts to test the suitability of the antibody reagents specified in the Registered Report (Bhargava et al., 2016), and reported in the original study (Hatzivassiliou et al., 2010) and observed that the monoclonal BRAF- and CRAF-specific antibodies did not appear to successfully detect the purified recombinant kinase domains (Figure 4A,B). To further test if the antibodies could detect the native, full-length target proteins, the antibodies were probed against whole cell extract from a panel of various human cell lines. This was an unexpected result, especially considering the data sheet from the supplier revealed a clear band by Western blot, however, the supplier result was from a nuclear envelope extraction of the HeLa S3 clone (Sigma Technical Service communication), while we tested a whole cell extraction from the parent HeLa cell line. While the CRAF antibody detected a signal with the appropriate molecular weight on SDS-PAGE gels, the BRAF antibody detected multiple bands (Figure 4C). In reviewing the original monoclonal antibodies that were used to detect BRAF and CRAF in the original study, it was evident that the immunogenic peptides used to generate these mouse antibodies included within their sequences about half of the amino acid residues that were not encompassed in the coding sequence for the recombinant BRAF and CRAF kinase domains (Figure 4D,E). The actual epitopes of these commercial antibodies were not determined and appear to be unknown. A possible explanation for these findings is that the recombinant BRAF and CRAF kinase domains we tested do not feature the immunoreactive epitopes, which can be as few as 5-10 amino acids. However, this requires further investigation, such as confirming the amino acid sequence of the recombinant kinase domains used as well as testing the sensitivity of the antibodies on full-length purified proteins.

To proceed with the dimerization assay, we identified different antibodies that successfully detected the recombinant BRAF and CRAF kinase domains (Figure 4A,B). During initial tests, we observed that the BRAF kinase domain immunoprecipitated with the CRAF kinase domain in the absence of any inhibitor, similar to the original study. However, the BRAF kinase domain also remained bound to Protein-A-agarose beads when the GST-CRAF kinase domain was not added to the assay (Figure 5A). This additional control was not conducted in the original study, but included in this replication attempt during optimization of the assay conditions and reagents. The positive detection of BRAF in this negative control condition, without the inclusion of GST-CRAF, revealed that the perceived binding of BRAF to CRAF, under these conditions, were largely, if not completely independent of the presence of CRAF. Instead, it appeared that the Protein-A-agarose bead suspension was binding to and pulling down the BRAF kinase domain, since this occurred in assays that did not include GST-CRAF or the anti-GST antibody (Figure 5B). This was also observed with different preparations of Protein-A-agarose beads. A further control that should be considered in the experimental design of future studies is the inclusion of the anti-GST antibody, to coat the Protein-A-agarose beads. Furthermore, wild-type BRAF appeared to bind much stronger to the Protein-A-agarose beads than the BRAF(V600E) mutant (Figure 5A,B). It is unclear how this compares to the original study, since the data represented in the original study does not directly compare the pull down of the two BRAF proteins.

**Figure 5.**
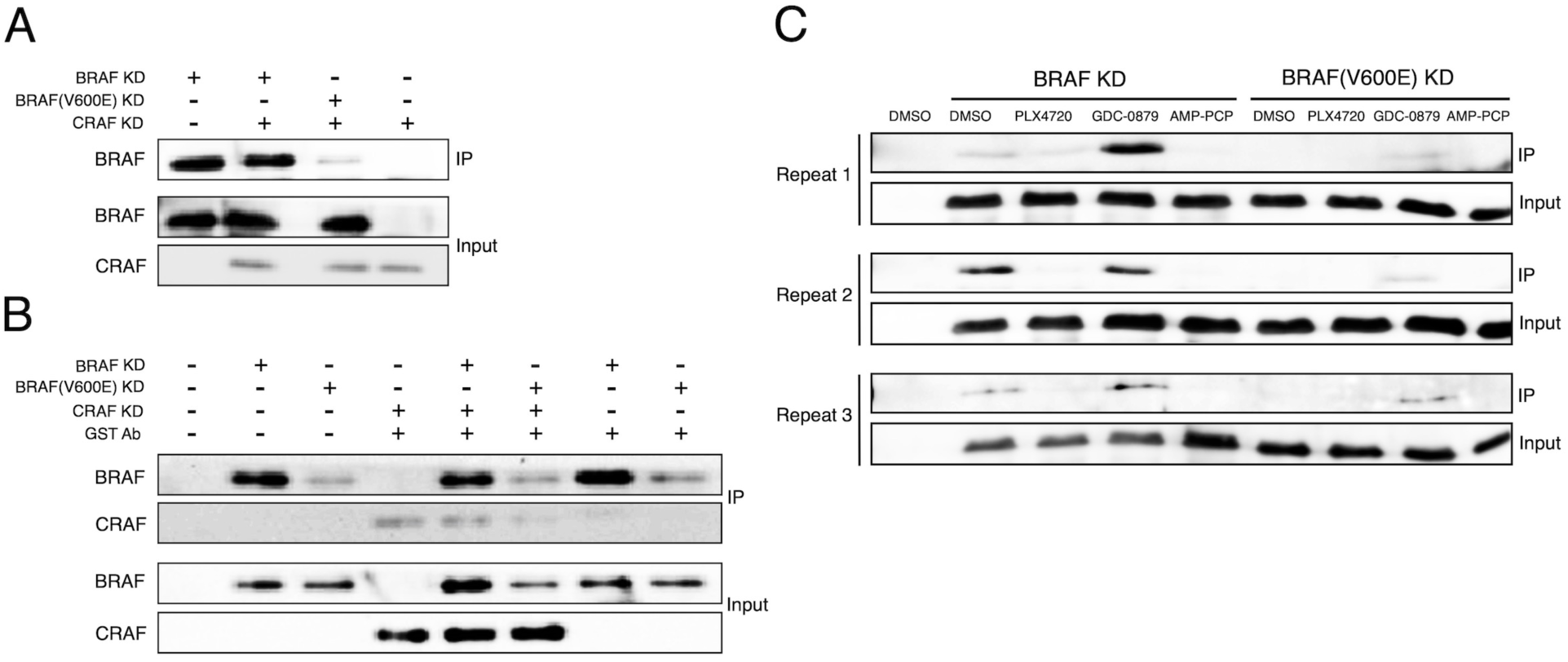
Biochemical dimerization assay with recombinant RAF proteins in the presence or absence of RAF inhibitors. Membranes were probed with BRAF (RRID:AB_2721113) or CRAF (RRID:AB_10989977) specific antibodies to detect BRAF and CRAF kinase domains, respectively. (**A**) Representative Western blots (experiment performed independently 2 times) of biochemical dimerization assay using purified recombinant GST-CRAF and BRAF kinase domains. IP performed with Protein-A-agarose beads and anti-GST antibody. (**B**) Representative Western blots (experiment performed independently 2 times) of biochemical dimerization assay using GST-CRAF and/or BRAF kinase domains. IP performed with Protein-A-agarose beads with or without anti-GST antibody. (**C**) Western blots of biochemical dimerization assay using BRAF kinase domains without GST-CRAF in the presence of DMSO, 10 µM PLX4720, 10 µM GDC-0879, or 1 mM AMP-PCP. IP performed with Protein-A-agarose beads without anti-GST antibody. Experiment performed independently 3 times, with each repeat shown. Additional details for this experiment can be found at https://osf.io/6f3sk/.

We examined whether the ATP-competitive RAF inhibitors differentially affected the direct binding of the recombinant BRAF kinase domain to the Protein-A-agarose beads. We found that the interaction between the wild-type BRAF kinase domain and the Protein-A-agarose beads were reduced in the presence of PLX4720 or AMP-PCP, and were increased in the presence GDC-0879 (Figure 5C,D). In contrast, the BRAF(V600E) kinase domain was not as strongly bound to the Protein-A-agarose beads and largely unaffected by these compounds. Although the intensity of the BRAF bands in the immunoprecipitates varied from experiment to experiment, the overall trends with respect to direct binding of wild-type BRAF to Protein-A-agarose, reduced binding with BRAF(V600E) and the various effects of the drugs were fairly consistent. A possible explanation for these findings is that the kinase inhibitors that were examined in this study, which bind to the active site of BRAF, likely produced conformational changes in the kinase domain structure selectively of wild-type BRAF and affected the overall stickiness of BRAF to protein and other surfaces. For example, 14-3-3 proteins, an essential cofactor for Raf kinase activity (Tzivion et al., 1998), have also been suggested to stabilize drug-induced BRAF-CRAF complexes (Garnett et al., 2005; Heidorn et al., 2010; Rushworth et al., 2006). Another non-mutually exclusive possibility is that GDC-0879, which has been reported to induce wild-type BRAF homodimers, could have resulted in greater amounts of BRAF on the Protein-A-agarose beads compared to PLX-4720, which has been reported to not stabilize BRAF dimers (Lavoie et al., 2013; Thevakumaran et al., 2015).

It is worth noting that while the GST-CRAF kinase domain used in this replication study had the same residue span (306 to 648) as the CRAF protein used in the original study, it was from a different supplier and the mutations to make CRAF constitutively active were changed from one negatively charged amino acid residue (original study: Y340D/Y341D) to another (this study: Y340E/Y341E), which might impact the results even though these changes are known to reproduce equivalent activations of the catalytic phosphotransferase activity of CRAF. Similarly, the recombinant BRAF kinase domain used in this replication attempt contained an additional 55 amino acids (417-766) compared to what was utilized in the original study (residues 432-726) (Figure 4D,E), which might impact the binding of the BRAF kinase domain to Protein-A-agarose beads. While it is unknown what impact these changes have in this *in vitro* biochemical dimerization assay, slight differences in the size of the BRAF and CRAF constructs used in previous studies do not appear to play a critical role in *in vivo* dimerization assays. For example, Jin et al. (2017) used catalytic domains of BRAF with residues 435-766 and CRAF with residues 327-648 to detect drug induced BRAF and CRAF binding by bioluminescence resonance energy transfer (BRET) in cells. An earlier BRET study from the same lab, Lavoie et al. (2013) used constructs with BRAF residues 444-723 and CRAF with residues 340-648. Thevakumaran et al. (2015) also used a shorter BRAF construct with residues 444-723 to explore the effects of drugs and mutations on BRAF dimerization in immunoprecipitation experiments.

Although not all the same reagents were utilized, as described above, we performed several optimization experiments in attempts to reduce the binding of recombinant BRAF to the Protein-A-agarose beads. These included extensive pre-washing of the Protein-A-agarose beads prior to incubation with the recombinant BRAF and CRAF kinase domains, using a different supplier of the Protein-A-agarose beads, producing our own agarose beads, substituting Protein-A beads with Protein-G beads, and using a different anti-GST antibody. Nevertheless, in all of these experiments, the BRAF kinase domain remained bound to the various antibody affinity bead preparations in the absence of added GST-CRAF and anti-GST antibody. Further, there was still a majority of the recombinant BRAF in the post-incubation supernatant fraction. It is unclear if these, or other technical aspects were problematic in the original study and if so, what was performed to address them. Additional approaches were also explored, such as using glutathione beads instead of the anti-GST antibody, which with further optimization might have made it possible to investigate the effects of the V600E mutation and impact of PLX4720, GDC-0879, and AMP-PCP on BRAF and CRAF heterodimerization. However, this was beyond the scope of this intended reproducibility study. Additionally, since so much optimization was required, the recombinant preparations of BRAF and CRAF may have deteriorated over time due to the extended time and multiple freeze-thaws while using and maintaining the proteins. This in addition to multiple batches of GST-CRAF that were needed to be utilized add additional factors that need to be considered when interpreting these results. These data, while unable to address whether, under the conditions of the original study, the same observations could be observed, highlight the challenges encountered during this replication attempt.

### Meta-analyses of original and replication effects

To compare the original and replication results for each experiment, where possible, the original reported value for each sample was compared to the distribution observed in this replication attempt. Some of the EC_50_ values between this replication attempt and the original study were not identical; however, overall they were consistent with each other. Specifically, the EC_50_ values for the RAF inhibitors, PLX4720 and GDC-0879, estimated from Figure 1A of Hatzivassiliou et al. (2010) were outside the 95% CI of the values for A375 cells observed in this replication attempt and were within the range of values observed in this replication for HCT116 and MeWo cells (Figure 1A,B). For the MEK inhibitor, PD0325901, the EC_50_ values estimated from Hatzivassiliou et al. (2010) were within the values observed in this replication for A375 and MeWo cells and outside the range of values for HCT116 cells (Figure 1C).

Similar to the EC_50_ values, the phospho-MEK levels determined after RAF inhibitor treatment in the original study were estimated *a priori* from the representative image reported in Figure 2B of Hatzivassiliou et al. (2010) and compared to the values reported in this replication attempt. For PLX4720, the phospho-MEK levels estimated from the original study were outside the 95% CI of the replication attempt, except the highest dose in the induced *CRAF* shRNA cells (Figure 3D). For GDC-0879, the phospho-MEK levels estimated from the original study were mixed (Figure 3F). For the *BRAF* shRNA conditions (with or without dox induction) the original values for the 0.1 µM and 1 µM doses were outside the 95% CI of the replication values, while the 10 µM dose was within the 95% CI. For the *CRAF* shRNA conditions, the estimated original values were all within the 95% CI of the replication attempt. Also, as stated above, in this replication attempt the activation of MEK by GDC-0879 was not the same between the *BRAF* and *CRAF* shRNA control cells. Importantly, the estimated values quantified from the images reported in Hatzivassiliou et al. (2010) should take into consideration that the images might not be in the linear dynamic range for measuring the relative expression of the target proteins (Bell, 2016; Taylor et al., 2013).

This direct replication provides an opportunity to understand the present evidence of these effects. Any known differences, including reagents and protocol differences, were identified prior to conducting the experimental work and described in the Registered Report (Bhargava et al., 2016). However, this is limited to what was obtainable from the original paper and through communication with the original authors, which means there might be particular features of the original experimental protocol that could be critical, but unidentified. So while some aspects, such as cell lines, inhibitors, and antibody manufacturers were largely maintained, others were changed at the time of study design, such as the length of the recombinant BRAF kinase domain, or during the replication attempt, such as the BRAF and CRAF antibodies, that could affect results. Furthermore, other aspects were unknown or not easily controlled for. These include variables such as cell line genetic drift (Hughes et al., 2007; Kleensang et al., 2016), differing compound potency resulting from different stock solutions (Kannt and Wieland, 2016), and lot to lot variability of reagents (Leek et al., 2010). Whether these or other factors influence the outcomes of this study is open to hypothesizing and further investigation, which is facilitated by direct replications and transparent reporting.

## Materials and Methods

As described in the Registered Report (Bhargava et al., 2016), we attempted a replication of the experiments reported in Figures 1A, 2B, and 4A of Hatzivassiliou et al. (2010). A detailed description of all protocols can be found in the Registered Report (Bhargava et al., 2016) and are described below with additional information not listed in the Registered Report, but needed during experimentation.

### Cell culture

A375 cells (ATCC, cat# CRL-1619, RRID:CVCL_0132) were grown in DMEM (ATCC, cat# 30-2002) supplemented with 10% Fetal Bovine Serum (FBS) (ATCC, cat# 30-2020). HCT116 cells (ATCC, cat# CCL-247, RRID:CVCL_0291) were grown in McCoy’s 5A (modified) medium (ATCC, cat# 30-2007) supplemented with 10% FBS. MeWo cells (ATCC, cat# HTB-65, RRID:CVCL_0445) were grown in EMEM (ATCC, cat# 30-2003) supplemented with 10% FBS. HCT116 cells with dox-inducible shRNA against *BRAF* or *CRAF* (Hoeflich et al., 2006) (shared by Genentech, Inc.) were grown in McCoy’s 5A (modified) medium (ATCC, cat# 30-2007 or Corning, cat# 10-050) supplemented with 10% FBS (Seradigm, cat# 1400-500G). Cells were grown at 37°C in a humidified atmosphere at 5% CO_2_. Quality control data for the A375, HCT116, and MeWo cell lines are available at https://osf.io/pma8b/, while the HCT116 cells with dox-inducible shRNA against *BRAF* or *CRAF* are available at https://osf.io/x42kr/. This includes results confirming the cell lines were free of mycoplasma contamination (DDC Medical, Fairfield, Ohio) as well as STR DNA profiling of the cell lines (DDC Medical), which were confirmed to be the indicated cell lines when queried against STR profile databases.

### Cellular dose response assay

The seeding density of each cell line was empirically determined by seeding at two-fold dilutions starting at 1.6×10^4^ cells per well in 96 well plates in 100 µl medium in technical quadruplicate. Cell viability was determined for the serially diluted cells 5 days after seeding with the Cell Titer Glo assay (Promega, cat# G7573) according to manufacturer’s instructions. Luminescence was quantified using a microplate fluorescence reader (BioTek, cat# FLx800). The number of cells seeded that remained sub-confluent by 5 days with a signal still in the exponential phase at the end of the assay were used to test the inhibitory effects of compounds.

MeWo cells were seeded at 1,500 cells/well, while A375 and HCT116 cells were seeded at 500 cells/well into 96-well plates and incubated overnight. The following day, cells were treated with serial dilutions of PLX4720 (Selleckchem, cat# S1152), PD0325901 (Selleckchem, cat# S1036), or GDC-0879 (Selleckchem, cat# S1104) to yield 9 dilutions of each drug ranging from 0.078 µM to 20 µM to give a final DMSO concentration of 0.1% v/v. Additionally, there were wells of untreated cells, vehicle control (0.1% v/v DMSO) treated cells, and media-only wells for each biological repeat. All conditions were done in technical quadruplicate. Wells were assayed for cell viability as described for the seeding density step 96 hr after start of treatment. For each biological repeat, the average background luminescence calculated from the media-only wells was subtracted from each treated well, thus the background luminescence from the media-only wells defined the baseline of 0% viability. Then the luminescence value of each treated well was normalized to the average of the vehicle control wells, thus defining the 100% viability as the average of the vehicle control wells. These data were fit to a four-parameter curve for each biological repeat to calculate the absolute half-maximum effective concentration (EC_50_) value for each drug treatment of each cell line as described in the Registered Report (Bhargava et al., 2016) using the *drc* package (Ritz et al., 2015) and R software (RRID:SCR_001905), version 3.5.1 (R Core Team, 2017). EC_50_ values unable to be accurately estimated following published guidelines (Sebaugh, 2011) were reported as >20 µM or <0.078 µM. Raw data files are available at https://osf.io/sb9xb/.

### Western blots from cells expressing inducible CRAF or BRAF shRNA

HCT116 cells expressing dox-inducible shRNA against *BRAF* or *CRAF* were seeded at 2.5×10^6^ cells/well in 6-well plates and treated with or without 2 µg/ml doxycycline. The concentration of doxycycline required to achieve a knockdown similar to what was reported in Figure 2B of Hatzivassiliou et al. (2010) was empirically determined by testing multiple doses of doxycycline (2 ng/ml, 20 ng/ml, 200 ng/ml, and 2 µg/ml) and comparing BRAF and CRAF expression levels to untreated cells as described in the Registered Report (Bhargava et al., 2016). After 72 hr, cells were treated with serial dilutions of PLX4720 (Symansis, cat# SY-PLX4720) or GDC-0879 (Selleckchem, cat# S1104) to yield dilutions of each drug (0 µM, 0.1 µM, 1.0 µM, and 10 µM) in a final DMSO concentration of 0.1% v/v. After 1 hr of treatment, total cell lysates were prepared by rinsing the cells in 1 ml ice cold PBS two times, lysing pellets in 0.4 ml per well of NP-40 Cell Lysis Buffer (Thermo Fisher Scientific, cat# FNN0021) containing Protease inhibitors (Roche, cat# 04693159001) and Phosphatase inhibitors (Thermo Fisher Scientific, cat# 78420). Cell lysates were clarified by centrifugation at 12,000x*g* for 5 min at 4°C. Protein concentrations were determined using a BCA Protein Assay (Thermo Fisher Scientific, cat# 23227) according to the manufacturer’s instructions. Samples concentrations were adjusted to 0.7 mg/ml, mixed with 2X Laemmli Buffer/ß-Mercaptoethanol (ß-ME), and stored at -20°C.

Samples were boiled for 5 min prior to loading 7 µg of samples on to 4-15% SDS-PAGE gels alongside a molecular weight ladder (Thermo Fisher Scientific, cat# LC5602) and electrophoresis at 200V for 40 min. Gels were transferred to nitrocellulose membranes using a Bio-Rad Trans-Blot Turbo Mini at 25 V, 2.5 A for 7 min. Completeness of protein transfer was determined by Ponceau S (Sigma-Aldrich, cat# P7170) staining for 5-6 min followed by frequent washing with H_2_0. Membranes were blocked for 1 hr with 5% w/v nonfat dry milk in 1X TBS with 0.1% Tween-20 (TBST). Membranes were probed overnight with rabbit anti-phospho-MEK1/2 (Ser217/221) (Cell Signaling Technology, cat# 9121; RRID:AB_331649) at 1:500 dilution in 1% BSA/TBST, rabbit anti-MEK1/2 (total MEK) [clone 47E6] (Cell Signaling Technology, cat# 9126; RRID:AB_331778) at 1:1000 dilution in 5% w/v nonfat dry milk/TBST, mouse anti-BRAF [clone F-7] conjugated to HRP (Santa Cruz Biotechnology, cat# sc-5284 HRP; RRID:AB_2721130) at 1:6000 dilution in 5% w/v nonfat dry milk/TBST, mouse anti-CRAF [clone 53/c-Raf-1] (BD Biosciences, cat# 610151; RRID:AB_397552) at 1:1000 dilution in 5% w/v nonfat dry milk/TBST, and mouse anti-ß-actin [clone 8H10D10] conjugated to HRP (Cell Signaling Technology, cat# 12262; RRID:AB_2566811) at 1:1000 dilution in 5% w/v nonfat dry milk/TBST. Following 3×5 min washes with TBST, membranes were incubated with the appropriate secondary antibody for 1 hr at room temperature: HRP-conjugated goat anti-rabbit (Thermo Fisher Scientific, cat# 31460; RRID:AB_228341) at 1:5000 dilution in 1% w/v BSA/TBST (or 5% w/v nonfat dry milk/TBST) or HRP-conjugated goat anti-mouse (Thermo Fisher Scientific, cat# 31432; RRID:AB_228302) at 1:5000 dilution in 5% w/v nonfat dry milk/TBST. Membranes were washed with TBST and incubated with ECL reagent (Bio-Rad, Hercules, California, cat# 1705060) according to the manufacturer’s instructions. Western blot images were acquired with ChemiDoc Touch Imaging System and Image Lab Touch software, version 1.1.0.04 (Bio-Rad) and quantified using the automated functions of Image Lab software (RRID:SCR_014210), version 5.2.1 (Bio-Rad). Additional details of image analysis are available at https://osf.io/zqmh4/. Additional methods and data, including full Western blot images, are available at https://osf.io/dqr5d/.

### Immunoprecipitation and Western blots for dimerization assay

N-terminal His6-BRAF and His6-BRAF(V600E) kinase domains were generated at the MRC Protein Phosphorylation and Ubiquitylation Unit (MRC PPU) Reagents and Services at the University of Dundee by expressing the baculovirus constructs pFBHTc delta N 1-416 BRAF [<u>DU586</u>] or pFBHTc delta N 1-416 BRAF V600E [<u>DU603</u>] in Sf21 insect cells at 27°C for 48 hr. Harvested cultures were affinity purified on Nickel-Agarose and stored in 50 mM Tris/HCl pH7.5, 0.1 mM EGTA, 150 mM NaCl, 0.1% ß-ME, 270 mM sucrose, 0.03% Brij-35, and then stored and shipped at -80°C. Purity was confirmed by resolving the purified kinases on a 4-12% gradient Bis-Tris protein gel and visualizing with a Coomassie-blue protein stain. The specific activity of the purified kinase domains were determined using the method described by Hastie et al. (2006). Initially, 5 µl His-BRAF protein (diluted in 50 mM Tris/HCl pH 7.5, 0.1 mM EGTA, 1 mg/ml BSA, 10 mM DTT) were incubated in a final volume of 20 µl of 50 mM Tris/HCl pH 7.5, 0.1 mM EGTA, 10 mM MgAc, 10 mM DTT, 1.4 µg ERK2, 0.4 µg MKK1, 0.1 mM ATP at 30°C for 30 min. Then, 2 µl of this reaction were assayed in a final volume of 50 µl of 50 mM Tris/HCl pH 7.5, 0.1 mM EGTA, 10 mM MgAc, 10 mM DTT, 0.33 mg/ml myelin basic protein, 0.1 mM [γ^32^P]ATP at 30°C for 10 min. Reactions were stopped by spotting 40 µl out of the 50 µl assay sample onto Whatman P81 papers, which were washed in 75 mM phosphoric acid, rinsed in acetone, dried, and counted. His-BRAF specific activity was 73192 U/mg (additional details are available at https://osf.io/47cqu/). His-BRAF(V600E) specific activity was 113858 U/mg (additional details are available at https://osf.io/2d8qp/).

The following kinase preparations were incubated in assay buffer (25 mM HEPES, pH 7.4, 10 mM MgCl_2_, 0.01% (v/v) Triton X-100, and 2 mM DTT) for 1 hr at room temperature: 500 nM (170 nM was used on some repeats to conserve material) of recombinant glutathione S-transferease (GST)-CRAF (Y340E/Y341E) kinase domain (sequence: 306-end) (SignalChem, cat# R01-13G); His-BRAF kinase domain (sequence: 417-end); and/or His-BRAF(V600E) kinase domain (sequence: 417-end) in the presence or absence of a fixed concentration of compound or vehicle (0.25% (v/v) DMSO). PLX4720 (Symansis, cat# SY-PLX4720), GDC-0879 (Selleckchem, cat# S1104), and Adenylylmethylenediphosphonate (AMP-PCP) disodium salt (Sigma-Aldrich, cat# M7510) were diluted in DMSO for their use. Proteins were immunoprecipitated with or without rabbit anti-GST antibody (Cell Signaling Technology, cat# 2622; RRID:AB_331670) at 1:56.6 dilution and Protein-A-agarose beads (Millipore, cat# IP02; also tested Kinexus, cat# BE-PRA-1). Complexes were centrifuged at 3000x*g* (HERMLE, Wehingen, Germany) for 10 min at 4°C to collect a sample of each fraction (e.g. supernatant) and to wash the pelleted beads several times with PBS by resuspension and centrifugation prior to resuspension of the pellet in SDS-PAGE sample buffer, which was boiled for 5 min to elute any proteins that may have bound to the beads. Samples alongside a molecular weight ladder were separated by SDS-PAGE (12.9% acrylamide gel) at 30 mA for 300 Vhr until the 10 kDa marker was at the bottom of the gel. Gels were transferred to nitrocellulose membranes at 250 mA for 1 hr and completeness of protein transfer was determined by Ponceau S staining. Membranes were blocked for 1 hr with 2.5% w/v nonfat dry milk and 1.5% BSA in 1X 0.05% TBST. Membranes were cut slightly below the 50 kDa marker to allow probing of BRAF and CRAF. Membranes were probed overnight at 4°C with mouse anti-BRAF [clone 3C6] (Sigma-Aldrich, cat# WH0000673M1; RRID:AB_1840330) at 2.5 µg/ml dilution, mouse anti-CRAF [clone 53/c-Raf-1] (BD Biosciences, cat# 610151; RRID:AB_397552) at 1:2500 (or 1:1000) dilution, rabbit anti-BRAF (Kinexus, cat# AB-NK156-4; RRID:AB_2721113) at 2 µg/ml dilution, or mouse anti-CRAF [clone H-8] (Santa Cruz Biotechnology, cat# sc-376142, RRID:AB_10989977) at 2 µg/ml dilution. Following washes with 0.05% TBST, membranes were incubated with the appropriate secondary antibody for 30 min at room temperature: HRP-conjugated donkey anti-mouse (Thermo Fisher Scientific, cat# SA1-100; RRID:AB_325993) at 1:500 dilution, HRP-conjugated goat anti-mouse (Santa Cruz Biotechnology, cat# sc-2005; RRID:AB_631736) at 1:5000 dilution, or HRP-conjugated donkey anti-rabbit (Santa Cruz Biotechnology, cat# sc-2077; RRID:AB_631745) at 1:10,000 dilution. Membranes were washed with 0.05% TBST and incubated with ECL reagent according to manufacturer’s instructions. Western blot images were acquired with Fluor-S Max Scanner (Bio-Rad) by scanning membranes for 320 sec with 40 sec intervals. Membranes were acquired with a Fluor-S MAX MultiImager (Bio-Rad) and Quantity One software (Bio-Rad, RRID:SCR_014280), version 4.1.1. Bands were quantified using ImageJ software (RRIS:SCR_003070), version 1.50a (Schneider et al., 2012). Additional methods and data, including full Western blot images, are available at https://osf.io/6f3sk/.

### Statistical analysis

Statistical analysis was performed with R software (RRID:SCR_001905), version 3.5.1 (R Core Team, 2017). All data, csv files, and analysis scripts are available on the OSF (https://osf.io/0hezb/). Confirmatory statistical analysis was preregistered (https://osf.io/rvnuc/) before the experimental work began as outlined in the Registered Report (Bhargava et al., 2016). Data were checked to ensure assumptions of statistical tests were met. When described in the results, the Bonferroni correction, to account for multiple testings, was applied to the alpha error or the *p*-value. The Bonferroni corrected value was determined by divided the uncorrected value (0.05) by the number of tests performed. Although the Bonferroni method is conservative, it was accounted for in the power calculations to ensure sample size was sufficient. The original study data was extracted *a priori* from the published figures by estimating the value reported. The extracted data was published in the Registered Report (Bhargava et al., 2016) and was used in the power calculations to determine the sample size for this study.

### Data availability

Additional detailed experimental notes, data, and analysis are available on OSF (RRID:SCR_003238) (https://osf.io/0hezb/; Pelech et al., 2018). This includes the R Markdown file (https://osf.io/k2xve/) that was used to compose this manuscript, which is a reproducible document linking the results in the article directly to the data and code that produced them (Hartgerink, 2017).

### Deviations from Registered Report

The lysis buffer to harvest HCT116 cells and source of PLX4720 and PD0325901 for the cellular dose response assay were different than what was listed in the Registered Report, with the used source and catalog number listed above. For the dimerization assay, the source of the GST-CRAF kinase domain was different than what was listed in the Registered Report, with source and catalog number listed above. The tag, size, and mutations were the same, except instead of Y340 and Y341 mutated to D, they were mutated to E, another negatively charged amino acid residue. Finally, as described in the results of the dimerization assay (Figure 5 & Figure 4), we also changed the source of the BRAF and CRAF antibodies used to detect the recombinant kinase domains by Western blot and modified the immunoprecipitation protocol when it was observed that the recombinant BRAF kinase domain was binding to the Protein-A-agarose beads independent of the CRAF kinase domain. Additional materials and instrumentation not listed in the Registered Report, but needed during experimentation are also listed.

## Acknowledgements

The Reproducibility Project: Cancer Biology would like to thank Shiva Malek (Genentech, Inc.) for sharing critical reagents and protocol information during preparation of the Registered Report, specifically the HCT116 cells with dox-inducible shRNA against *BRAF* or *CRAF*. We would also like to thank the MRC Protein Phosphorylation and Ubiquitylation Unit (MRC PPU) Reagents and Services at the University of Dundee for preparing the recombinant BRAF and BRAF(V600E) kinase domains, Sara Marrello for assistance with the dimerization assay, and the following companies for generously donating reagents to the Reproducibility Project: Cancer Biology; American Type and Tissue Collection (ATCC), Applied Biological Materials, BioLegend, Charles River Laboratories, Corning Incorporated, DDC Medical, EMD Millipore, Harlan Laboratories, LI-COR Biosciences, Mirus Bio, Novus Biologicals, Sigma-Aldrich, and System Biosciences (SBI).

## Competing Interests

SP, CS, LY: Kinexus Bioinformatics Corporation is a Science Exchange associated lab. SP and CS hold the majority of the shares in Kinexus.

CG, JK: University of Maryland College Park is a Science Exchange associated lab.

AB: Shakti BioResearch LLC is a Science Exchange associated lab.

EI, RT, NP: Are, or were, employed by and hold shares in Science Exchange Inc.

T.M.E., A.D.: Are, or were, employed by the nonprofit Center for Open Science that has a mission to increase openness, integrity, and reproducibility of research.

